# Clonal analysis reveals gradual recruitment of lateral root founder cells and a link between root initiation and cambium formation in *Arabidopsis thaliana*

**DOI:** 10.1101/283366

**Authors:** Joseph G. Dubrovsky

## Abstract

- The pericycle gives rise to lateral roots (LRs) and lateral meristems (LMs; cambium and phellogen), however, a thorough clonal analysis of pericycle cell lineage has not been investigated. This study fills in this gap and addresses pericycle impact in LR and LM development.
- Heath-shock inducible *DS1* transposition in *35S-DS1-H2B:YFP; HS-Ac* seedlings results in production of YFP-labelled cell clones. These clones in pericycle cell derivatives were identified with a confocal microscopy and subjected to 3D reconstructions and analysis.
- Participation of pericycle founder cells (FC) in LR formation is more variable than previously considered. LR initiation was found most commonly involved the specification of just one FC in the longitudinal and one or two cells in transverse direction. After LR initiation, FCs continue to be recruited in both directions from pre-existing cells. Anticlinal divisions in the pericycle resulting in LMs start already in the young differentiation zone where only the protoxylem is differentiated.
- The clonal analysis demonstrated that pericycle cell activity related to LR formation is not separated in time and space from that related to LM formation and that LR FC recruitment is a gradual process. The analysis demonstrated that immediate pericycle progeny lack self-renewal capacity.

## Introduction

In plants, contrary to animals, during post-embryonic development, organogenesis is maintained during lifespan of an organism due to activities of the apical and lateral meristems (LMs) which permit organ growth in length and girth, respectively. New apical meristems are continuously produced. A single plant is capable to produce an order of 107 of lateral roots (LRs) for four months (Dittmer, 1937). This process takes place in a specified tissue, pericycle. In *Arabidospis thaliana*, pericycle is a one-cell-layer thick and the external most tissue of vascular (central) cylinder or stele, which surrounds xylem, phloem, and vascular parenchyma cells (Fig. 1a). Xylem pole pericycle (XPP) cells continue cycling after leaving the root apical meristem (Dubrovsky et al., 2000, Beeckman et al., 2001) but the competence of pericycle cells to form LRs is restricted in time and space and takes place at certain developmental window (Dubrovsky et al., 2006, 2011). After this window, pericycle still maintains its proliferation capacity and forms two new LMs, cambium and cork cambium (phellogen). In spite of wide use of this model organism, we still lack the complete understanding of pericycle contribution to LR and LM formation and do not know whether these activities are coupled.

**Fig. 1.**
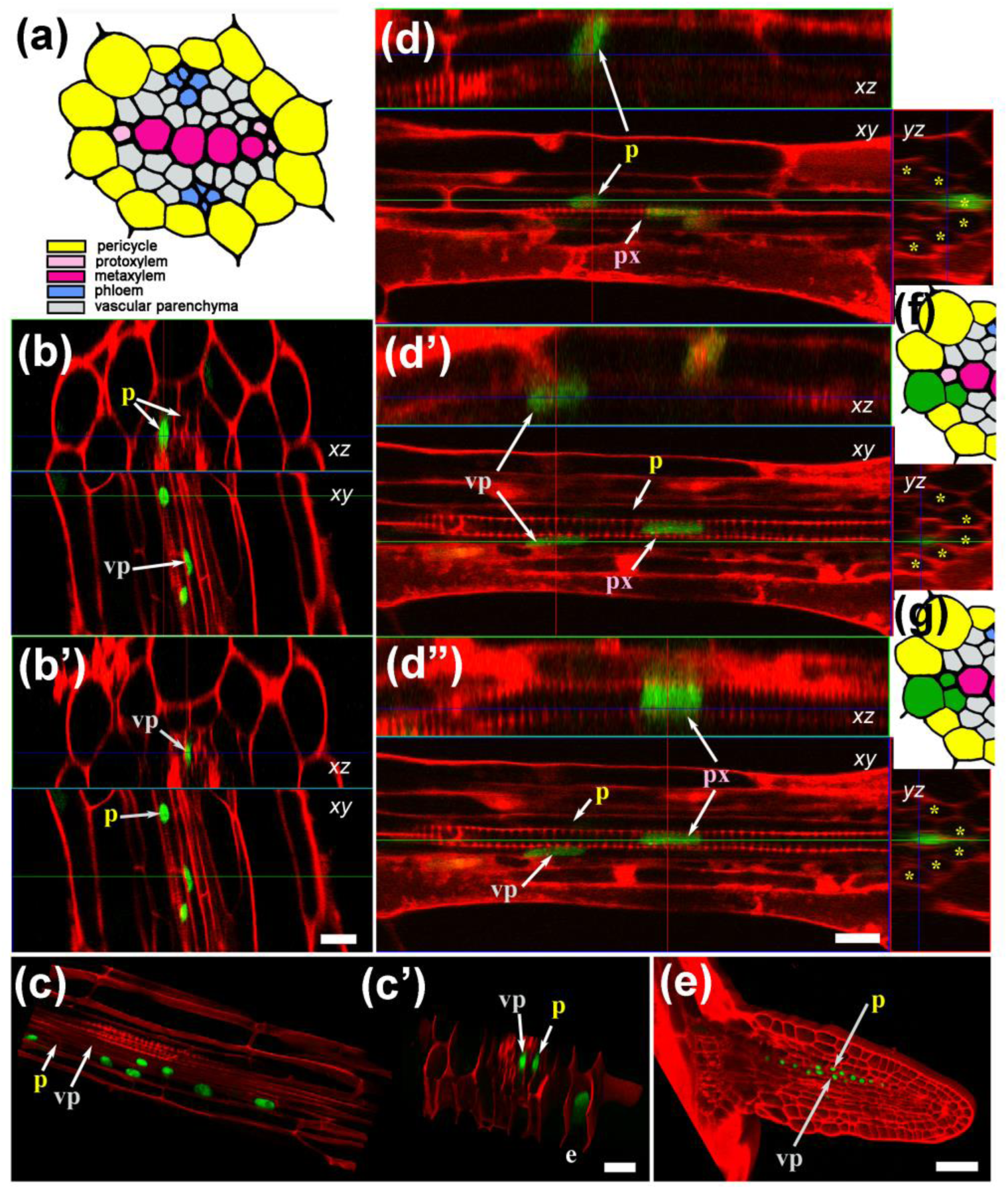
Clonal relationships of the pericycle and other cell types of the stele. (a) Schematic of the primary tissues of the Arabidopsis root stele in the transverse plane. The external layers (endodermis, cortex and epidermis) are not shown for simplicity. (b, b’) A clone of H2B-YFP-marked cells of the pericycle and vascular parenchyma. (c, c’) A 3D reconstruction of a common clone of marked cells containing the pericycle and a dividing vascular parenchyma cell (shown in c). (d–d’’) A clone comprising one xylem pole pericycle (XPP) cell, a vascular parenchyma cell, and a differentiating protoxylem cell. (e) A recently emerged lateral root, in which a clone of the pericycle and vascular parenchyma cells has formed. (f) The clonal relationship of XPP and vascular parenchyma cells (green) corresponding to the clones shown in (b), (c) and (e). (g) The clonal relationship of XPP, the vascular parenchyma and protoxylem cells (green), corresponding to the clone shown in (d). Panels labelled with the same letter depict the same root image taken at different x-y-z positions. Panels of images taken in xy, xz and yz sections are indicated. p, pericycle; vp, vascular parenchyma; e, epidermis; px, protoxylem; asterisks, pericycle cell layer. Scale bars: 10 µm (b), 20 µm (c, d) and 40 µm (e).

During LR formation, a plant hormone auxin acts as a morphogenetic trigger and specifies certain pericycle cells as founder cells (FCs) for LR primordium (LRP) formation (Dubrovsky et al., 2008). Longitudinal bi-cellular type of initiation comprises two longitudinally adjacent FCs in a cell file give rise to entire primordium in the longitudinal plane. It has been identified in a number of species (Casero et al., 1995) and in *Arabidopsis* it is common to consider that this is the only type of LR initiation which was well documented in anatomical and time-lapse studies (Casimiro et al., 2001; Dubrovsky et al., 2001, 2008; De Smet et al., 2006, 2008; De Smet, 2012; De Rybel, et al., 2010; Lucas et al., 2013; Jansen et al., 2013; Fernandez et al., 2015). However, another type, longitudinal uni-cellular, when a single FC in a cell file gives rise to entire LRP longitudinal extension, has also been recognised (Dubrovsky et al., 2001, 2008). Until now, however, we do not know how prevalent are these types of initiation in *Arabidopsis* and whether there are some other type(s) of LRP initiation. Importantly, even time-lapse cell tracking studies do not permit to answer this question (von Wangenheim et al, 2016). It is commonly considered that in transverse plane three contiguous XPP are involved in initiation of the LRP (Dubrovsky et al., 2001; Casimiro et al., 2003; Parizot et al., 2008). Serial histological sectioning showed that up to four XPP cells can be activated during LRP initiation (Dubrovsky et al., 2001). Recently it was found that five to seven cell files of contiguous XPP cells become activated (von Wangenheim et al, 2016). Here, clonal analysis has been performed to clarify what scenario of pericycle cells participation in LR formation is most common.

Clonal analysis permits understanding the morphogenetic components of organ development, such as distribution, orientation and abundance of cell divisions, cell fate, and estimation of the number of cells giving rise to an organ (Poethig, 1987; Buckingham and Meilhac, 2011). However, thorough clonal analysis of LR formation is laborious and last time was performed more than a half a century ago (Davidson, 1965). In this study, pericycle cell lineage was investigated and it was found that pericycle contribution to LR formation is gradual and coupled to pericycle participation in LM development and that these pericycle activities start already in the young differentiation zone. The data indicate that the pattern of pericycle cell division related to LR initiation is more variable than generally considered and that some dividing pericycle cells and vascular parenchyma or xylem cells can share common progenitors.

## Materials and Methods

### Plant material and growth conditions

The heat-shock inducible, Activator (Ac)-Dissociator (Ds) lineage marking technique used in this study was described (Kurup et al., 2005). Briefly, the authors constructed two lines, both in Landsberg erecta (Ler), one carrying heat shock inducible promoter fused to Ac transposase (HS-Ac) and another with 35S promoter fused to *H2B:YFP* and interrupted with Ds1 transposable element (*35S-DS1-H2B:YFP*). In the latter line, Ds1 interrupts YFP reporter gene expression. After a 42o C heat shock, Ds1 is excised by Ac transposase and a single cell where transposition took place shows nuclear YFP expression. The progeny of the marked cell forms a clone (Kurup et al., 2005). These authors reported that some F1 progenies showed a high score of transposition events whereas others had much lower score or did not show transposition events. Parent lines and few F1 crosses were kindly donated to us by Drs. Smita Kurup (Rothamsted Research, Hertfordshire, UK) and Laurent Laplaze (Institute of Research for Development, Marceille, France). Seven lines were used with the highest score of transpositions and next generations were obtained. F2 generation of line 7 seedlings were subjected to 42o C heat shock and analysed under an epifluorescent microscope. Those individual plants which showed one or more transposition events were carefully transferred to soil and their F3 and F4 progeny (individual lines) subsequently were used for this study. New crosses were made between the *HS-Ac* and *35S-DS1-H2B:YFP* lines and F1 plants were used for clonal analysis. Overall 17 lines were used for clonal analysis. These lines produced similar clones of YFP-marked cells and, irrespectively of generation, were used for analysis. Different heat shock durations were tested (ranging from 20 to 180 minutes) and 45 min was found to be optimal. In most cases the plants were analysed during the next day of growth after a heat shock treatment. YFP-NPSN12 plasmalemma marker line *Wave 131YFP* (Geldner et al., 2009), *35S:H2B:RFP* line (Federici et al., 2012), and *pGATA23::NLS-GFP* (De Rybel et al., 2010), *alf4-1* mutant (DiDonato et al., 2004) and *alf4-1;DR5rev:GFP* (Dubrovsky et al., 2008), all in Col-0 background, were described. Each individual *alf4-1;DR5rev:GFP* seedling was genotyped with PCR as described (Dubrovsky et al., 2008) and only roots of homozygous *alf4-1* seedlings were subjected to confocal laser scanning microscopy (CLSM) analysis. For root growth dynamics, number of replicates and statistical test are indicated in the respective figure legend (Fig.S1).

Seeds were surface sterilized and grown in 0.5× MS salts, 1% sucrose, 0.8% agar in a growth room at 21o C and 16h light photoperiod. In most cases, heat shock was performed on plants five days after germination. Prior CLSM seedlings were analysed under a Leica MZFL III binocular microscope equipped with epifluorescent system and appropriate YFP settings (Leica Microsystems, Milton Keynes, UK).

### Microscopy

Live roots were stained with 5 μg/ml of propidium iodide dissolved in water. CLSM studies were performed with a Zeiss LSM 510 Meta (Oberkochen, Germany) microscope using 488 nm line of Argon laser, yellow channel emission filter BP 505-550 nm and a secondary dichroic filter NFT 635 nm. Zeiss C-Apochromat 40x/1.2 W corr objective was used in most cases. Some experiments were performed on a Leica SP5 CLSM (Leica Microsystems, Milton Keynes, UK). Maximum or average intensity projections were obtained with Image J (http://rsb.info.nih.gov/ij), or LSM browser which also was used for 3-D reconstructions. MorphoGraphX (www.MorphoGraphX.org, de Reuille et al., 2015) was used for virtual sectioning and reconstructions. Technical details of the illustrations presented in the article are specified in Table S1.

### Clonal analysis

In total, 52 experiments were performed. Total number of seedlings analysed for root clones was >1192. Individual plants that formed YFP-expressing sectors were marked and 218 seedling were analysed under a CLSM. Under the growth conditions, the incidence of transposition events was low and only clones formed in pericycle or its derivatives were included in the analysis. By these reasons, no cell tracking of clones was performed. In total 136 clones were identified (Table 1). Information about size of clones, root growth in *HS-Ac;35S-DS1-H2B:YFP* seedlings before and after heat shock can be found in Supporting information online.

**Table 1.**
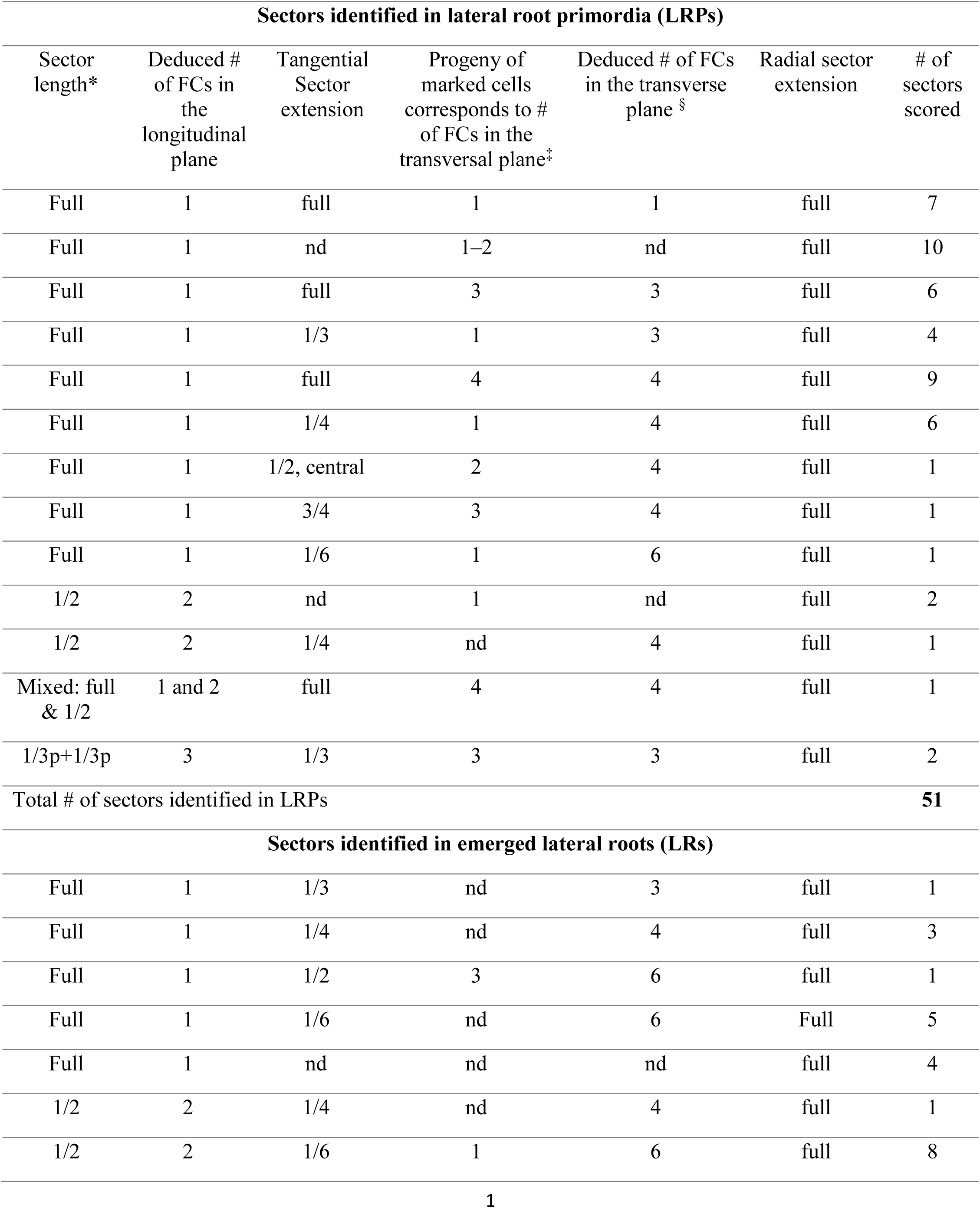
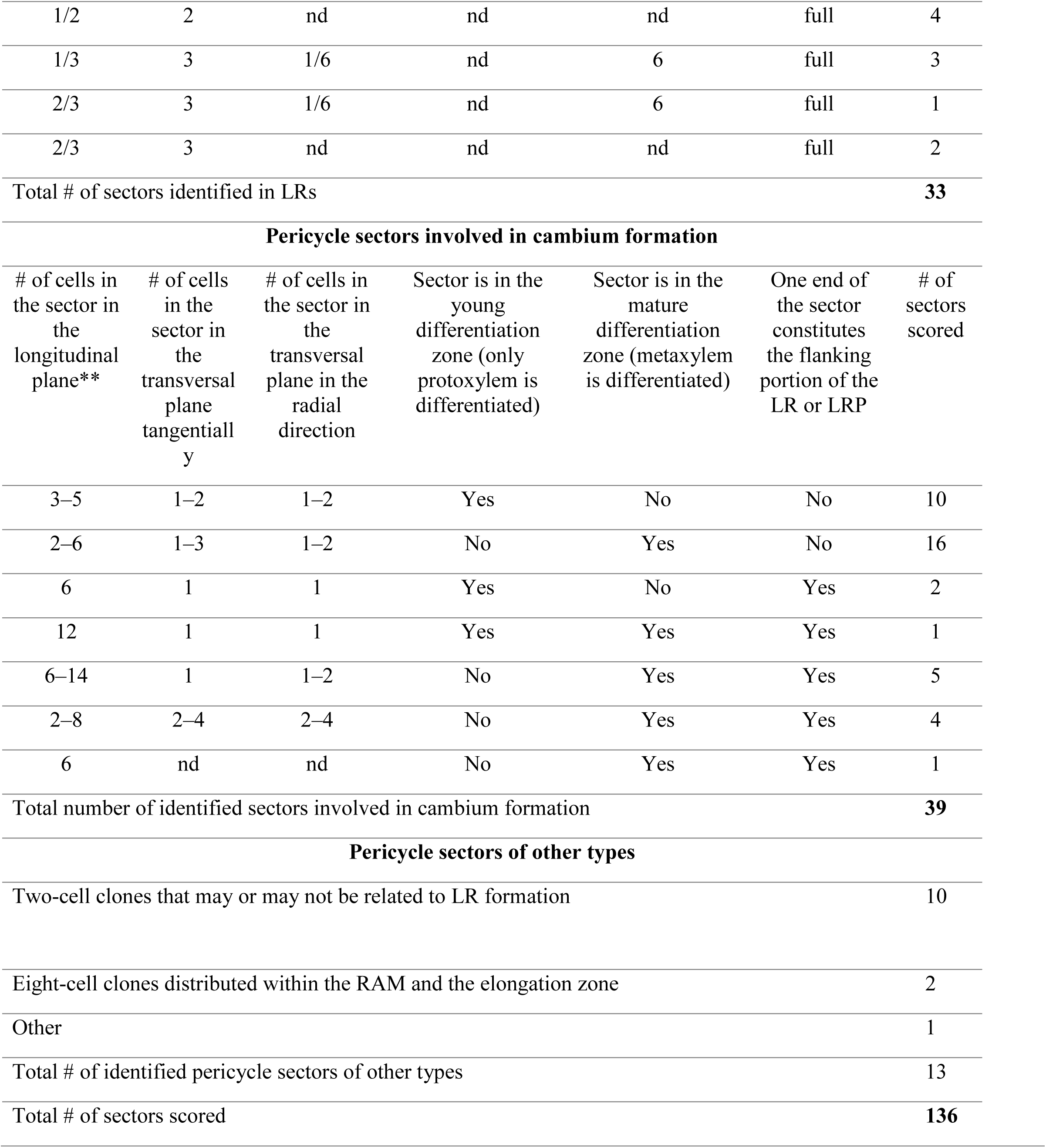
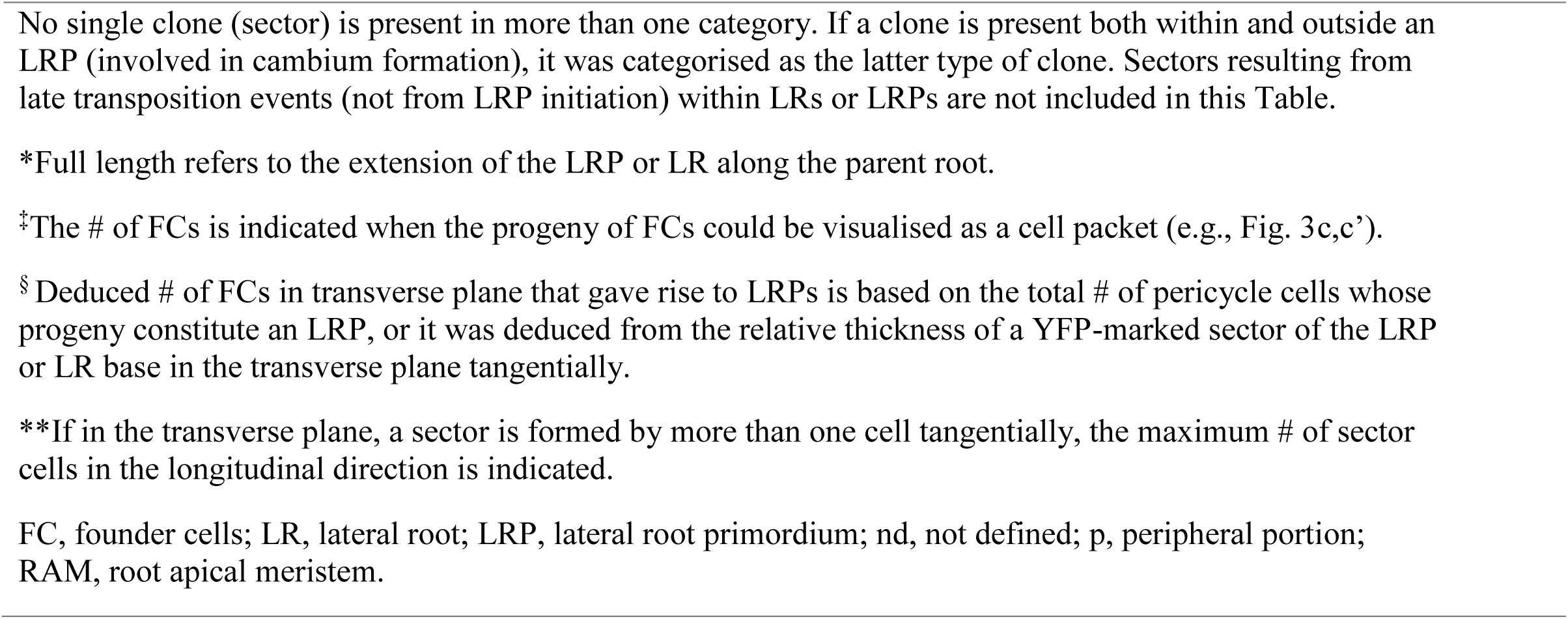
Classification of clones (sectors) identified in the pericycle or its derivatives and their number (#)

## Results

Pericycle cell lineage in the *A. thaliana* root was studied using *35S-DS1-H2B:YFP; HS-Ac* seedlings subjected to heath-shock leading to transposition of a DS1 transposon and resulting in production of YFP-labelled cells (Kurup et al., 2005). A clone was formed as a result of a division of these cells. Single differentiated cells were also subjected to the transposition suggesting that sectors were not formed preferably in dividing cells. Root growth dynamics of *35S-DS1-H2B:YFP; HS-Ac* seedlings was comparable with those in wild type (Fig. S1). The terms ‘clones’ and ‘sectors’ are used as synonyms. The terms ‘longitudinal’ and ‘transverse’ refer to the respective planes within the primary root. Anatomical coordinate terminology used in this study is illustrated on Fig. S2.

### Protoxylem, vascular parenchyma, and pericycle can share common progenitors

The pericycle gives rise to LRs and participates in formation of vascular and cork cambia that produce secondary tissues (Dolan et al., 1993). It is not known whether pericycle can share common origin with vascular parenchyma (Fig. 1a), also termed ‘procambium’ (Mähönen et al., 2000). Total number of pericycle-related clones found was 136 (Table 1). Four clones (2.9%) were found in which pericycle and other cells types of the stele originated from a common progenitor cell. Some clones included two pericycle cells and a single vascular parenchyma cell (a three-cellular clone depicted on Fig. 1b) or five pericycle cells and a vascular parenchyma cell that was dividing (Fig 1c). These clones indicate that a common progenitor cell gave rise to pericycle and vascular parenchyma cells (Fig. 1f). In a clone depicted on Fig. 1c pericycle nuclei are closely spaced suggesting that anticlinal divisions took place after FC cell left the meristem. Longitudinal size of the group of YFP-marked cells in the pericycle suggests that this is a stage I LRP. Mother cell of this pericycle clone and a mother cell of a dividing vascular parenchyma cell probably formed a common clone already in the apical meristem and thus share a common origin. A vascular parenchyma cell was in telophase stage of mitosis indicating that vascular parenchyma cells are capable to maintain proliferation activity outside the root apical meristem. The clone depicted in Fig. 1d-d’’ shows YFP-marked adjacent XPP, vascular parenchyma, and protoxylem cells, one of each type. This indicates that a single progenitor cell gave rise to these tree cell types of the central cylinder (Fig. 1g). The fact that a YFP-marked nucleus of theprotoxylem cell was found in the young differentiation zone suggests that the clone was formed before the exit of cells from the root apical meristem.. Also, in accordance with geometry of the location of these cell types on the transversal section (Fig 1g) one can suggest that a progenitor cell divided periclinally twice: once in tangential and once in radial directions.

No pericycle-related clones were found within the apical meristem of the primary roots. However, one clone was found in the recently emerged LR that contained YFP-marked cells in the pericycle and the adjacent internal layer (Fig. 1e). Cells in the rootward portion of this clone maintained meristematic features. This clone thus showed the possibility that a common progenitor cell within the apical meristem gives rise to pericycle and a more internal cell type. The probability of two independent transposition events in the pericycle and a neighbouring tissue is lower than that of a single transposition event. Therefore, the clones of YFP-marked cells that include pericycle, the protoxylem, and vascular parenchyma indicate that these cell types can share common progenitors. These clones also confirm that position and not lineage defines the cell fate (Kidner et al., 2000).

### Participation of pericycle founder cells in lateral root formation is more variable than previously considered

In *Arabidopsis*, an LRP is initiated in pericycle at the distance from the root tip where protoxylem cells are already differentiated. To determine the LRP progenitor cell number, clones were analysed in longitudinal and transverse planes.

#### Longitudinally one to three pericycle cells can give rise to a primordium

To establish the number of progenitor cells giving rise to an LRP, it is important to consider what proportion of the developing LRP or LR forms a clone (sector) and what is the timing of origin of each clone. When only a fraction of developing LR contained YFP-marked cells, the timing of origin of the clones can be variable and not necessarily linked to the LR initiation. Therefore, clones spread at the base of the LRP or LR to whole longitudinal extension, were formed from the LR initiation. It was found that the clones corresponding to only one progenitor cell in the longitudinal direction were most common. Further the types of the clones found are described.

In the young differentiation zone, defined as the distal root region where metaxylem is not yet differentiated, there were clones with a single YFP-marked XPP cell (Fig. 2a) or two closely spaced marked cells in one cell layer (Fig. 2b). I also found one-cell-layer thick clones of short pericycle cells similar to a five-cellular clone of YFP-marked cells depicted on Fig. 2c. All these clones are stage I LRP (classification of Malamy and Benfey, 1997). If longitudinally only one FC participates in primordium formation, all the cells along the base of the prospective LR should be YFP-marked (Fig. 2b-f). Shootward and rootward of the described clones no other YFP-marked pericycle cells or their derivatives were found. As the clones were detected 24 h after heat shock, and time from cell displacement form the apical meristem till the first LRP stages is 13.6 to 16 h (Dubrovsky et al., 2000) and some time is required for DS1 excision and GFP maturation, the data suggest that the cell that became recruited later as the FC was subjected to an excision event already in the root apical meristem. Then this type of clones suggests that only one FC gave rise to the entire longitudinal extension of the developing LRP (Fig. 2g) supporting the scenario that a single cell in the longitudinal orientation acts as a FC of the LR (Dubrovsky et al., 2001).

**Fig. 2.**
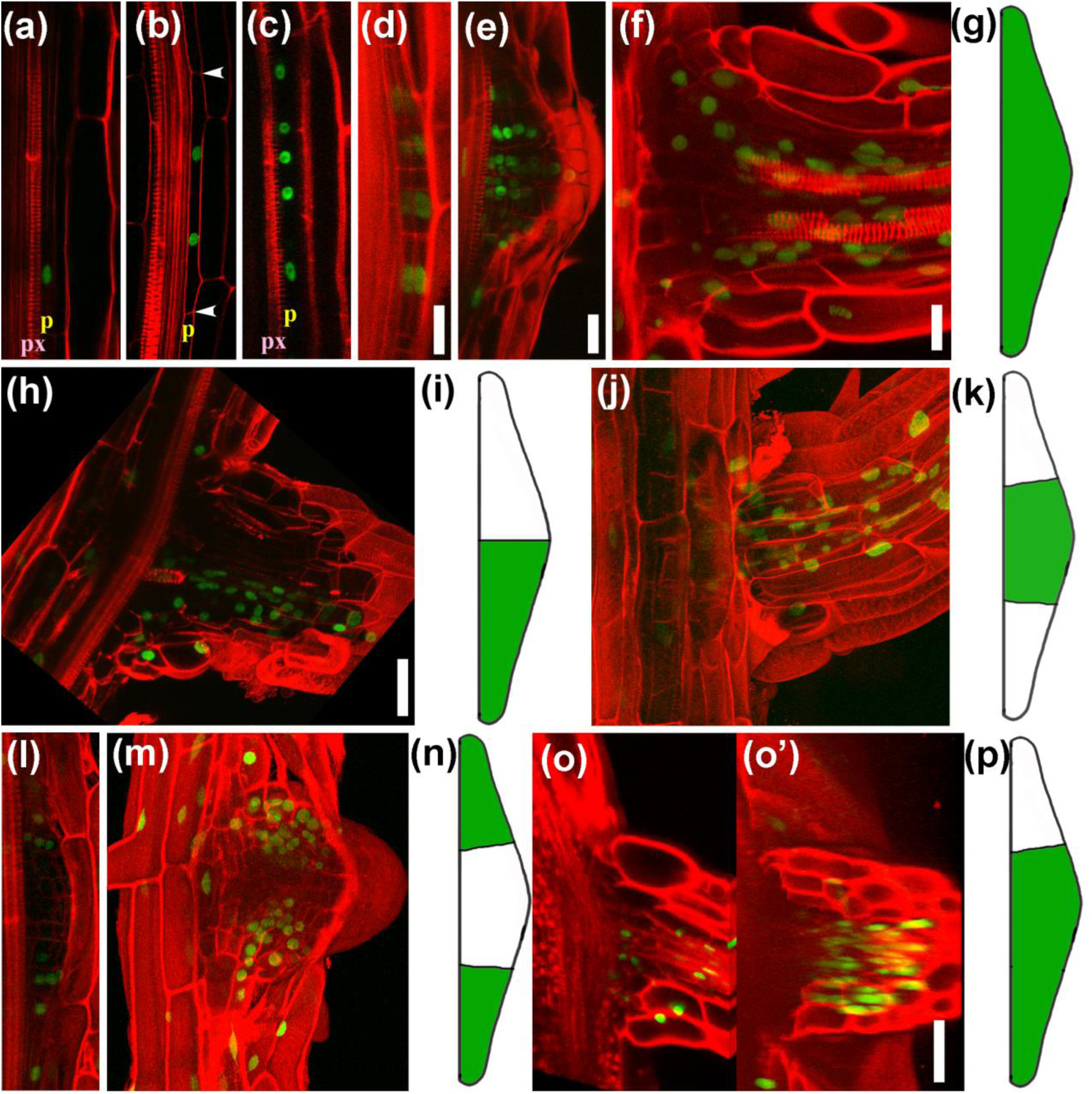
Clones analysed in the longitudinal plane of the parent root. (a–f) Clones that correspond to longitudinally unicellular lateral root primordium (LRP) initiation. (a) A single marked xylem pole pericycle cell. (b) A bicellular longitudinal clone of a stage 0 LRP. The borders of the clone are indicated with arrowheads. (c) A clone distributed longitudinally along the entire length of stage I LRP. (d–f) A stage II LRP (d), a stage V LRP (e) and an emerged LR (f) with clones distributed along the entire thickness of the organ in the longitudinal plane. (g) Schematic depiction of the clones corresponding to one cell in the longitudinal plane giving rise to an LRP or LR. (h, i) A clone distributed along half of the thickness of an emerged LR. (i) Schematic depiction of the clones in (h). (j) A clone distributed along one-third of the thickness of the emerged LR. (k) Schematic depiction of the clone in (j). (l and m) A stage IV LRP (l) and a recently emerged LR (m) containing clones of marked cells at their flanks, occupying approximately one-third of the organ thickness in the longitudinal plane. (o) A longitudinal section and (o’) a 3D reconstruction of an LR with a clone distributed across approximately two-thirds of its thickness in the longitudinal plane. (p) Schematic depiction of the marked sector of the root shown in (o). Scale bars: 40 µm. Scale bar in (d) corresponds to (a–d). Scale bar in (o’) corresponds to (j), (l), (m), and (o).

Clones that ran approximately half the length of the LR thickness (in the longitudinal direction of the parent root) were also found (Fig 2h). In the other half (upper portion of the panel 2h) there was just a single YFP-marked nucleus indicating a late transposition event. This indicates that two progenitor pericycle cells in longitudinal direction could give rise to the LR and this type then should correspond to bi-cellular longitudinal type of LRP initiation (Dubrovsky et al., 2001). Another possibility is that this type of clones resulted from a division of a single cell (in the longitudinal orientation; outline on Fig. 1g) but *Ds* transposition occurred later and only in one of the daughter cells. Then, this type of clones does not exclude that they belong also to uni-cellular longitudinal type of LRP initiation.

Another type of longitudinal clones was identified when central 1/3 portion of the LR thickness in the longitudinal direction of the parent root was marked (Fig 2j, k). This type then should correspond to two models: (1) model where three progenitor cells in the longitudinal direction give rise to LR and (2) model where *Ds* transposition occurred not at initiation time but later. Then this type of clones can correspond to longitudinally one or two progenitor cells. Some LRs and LRPs had approximately 1/3 of LR or LRP (central portion) not marked but approximately 2/3 of their thickness in longitudinal direction (1/3 located rootward and 1/3 shootward of the central portion) was marked (Fig. 2l,m,n). These clones support the same two models outlined above. Some intermediate scenario was also identified. In some LRs, 2/3 of the root thickness in the longitudinal direction was marked (Fig. 2o,p). This type could correspond either to unequal participation of two progenitor cells or approximately equal participation of three progenitor pericycle cells in the longitudinal direction. The most plausible interpretation of this clone is that the 2/3 of LR extent in the longitudinal plane was formed from one progenitor cells and the rest of the LR from another one (Fig. 2o,o’) or this was a later event after a longitudinally single progenitor cell gave rise to the LRP.

If to consider only the cases depicted on Fig. 2o,p, as of bi-cellular type of initiation, then out of total 84 clones in LRP and LR, 59 (70%) were of uni-cellular type, 20 (24%) were of bi-cellular type and 5 (6%) were of the type when longitudinally three FCs cells could give rise to an LRP (Table 1). This suggests that overall longitudinal uni-cellular type of initiation was more common.

#### Analysis of clones in transverse plane indicates that lateral root founder cell recruitment continues after lateral root primordium initiation

To estimate the number of pericycle cells participating in LR formation in the transverse plane the size and shape of clones in advanced LRP stages (stage IV and later) and in developed LRs are analysed first and then in early stage LRPs. This analysis suggests that in these two age groups the number of FCs participating in LR formation is different and so it should be increasing from young to later LRP stages.

Analysing clones at advanced LRP stages and in LRs, I found different types of clones outlined in Fig. 3. In some cases, a clone occupied approximately half of the LRP in transverse plane (Fig. 3a,b). In this LRP, based on anatomical cell arrangements, the YFP-marked portion corresponded to three sub-sectors (cell packets depicted by yellow circles) of similar size similar to other three sub-sectors of unmarked cells (Fig. 3a). Each cell packet contains the progeny of a a FCs as reported in a time-lapse studies (Wangenheim et al., 2016, Goh et al., 2016). This analysis suggests that there were in total six progenitor cells in transverse plane that gave rise to the LRP. Other LRPs were also identified in which one clone in transverse plane occupied position of a single sub-sector (cell packet) of YFP-marked cells (Fig. 3c,c’,d). In this LRP, the clone was about 1/4 of the transversal area of the primordium in which in total four sub-sectors in transverse plane were detected (Fig. 3c,c’,d). This suggested that four progenitor cells gave rise to the primordium. Analysing emerged LRs when viewed in the transverse plane, I found roots with clones of YFP-marked cells that were spread to approximately 1/6 of the LR thickness. These clones were formed in central (Fig. 3e and Fig.S2) or peripheral (Fig. 3g-h) sub-sectors of the developing LRP. These cases suggested that in transverse plane six progenitor cells of the parent root can give rise to the LR, a finding corresponding well with that of Wangenheim et al. (2016).

**Fig. 3.**
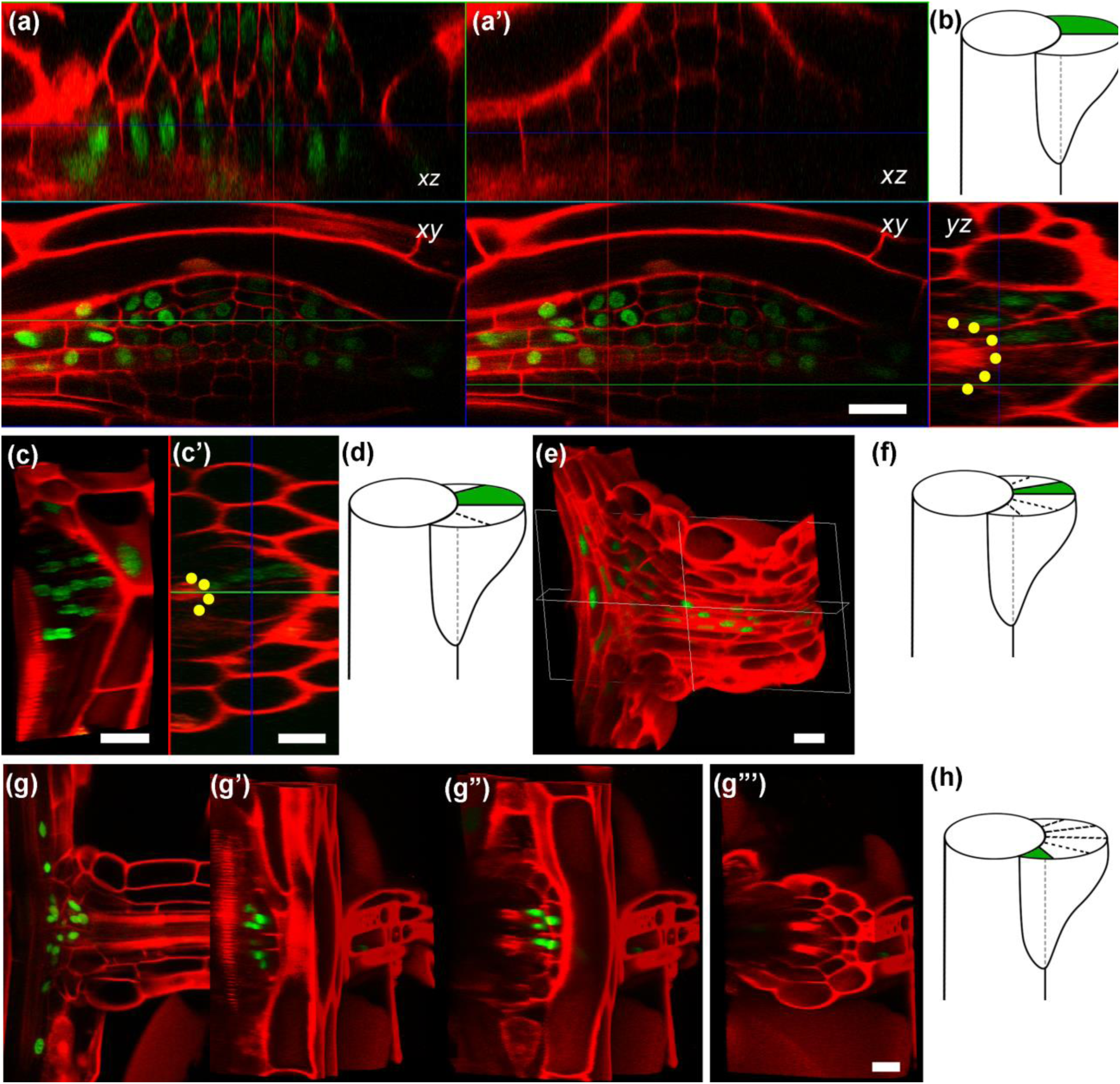
Clones analysed in the transverse plane of the parent root. (a) A lateral root primordium (LRP) containing a clone that comprises about half of its transverse plane. (b) Schematic depiction of (a). (c and c’) A 3D reconstruction showing a sector of marked cells corresponding to one-quarter of the LRP in the transverse plane. (d) Schematic depiction of (c). (e) A 3D reconstruction of an LR containing a clone of marked cells about one-sixth of its thickness in the transverse plane, and a corresponding scheme (f); see also Fig. S3. (g) A longitudinal optical section of a recently emerged LR containing a clone of marked cells developed at the periphery of the LR base. (g’–g’’’) 3D reconstructions of the same root showing the distribution of a marked clone across one-sixth of the lateral root base in the transverse plane, located at its periphery. (h) Schematic depiction of the clone shown in (g–g’’’). Yellow dots in (a’) and (c’) indicate the position of the FCs in the transverse plane. The axes xy, xz and yz are as they were in Fig. 1. Scale bars: 20 µm in (a), (c), (e) and (g–g’’’); 10 µm in (c’).

During LRP initiation, a subset of long pericycle FCs divide anticlinally and give rise to stage I LRP composed of short pericycle derivative cells arranged in one cell layer. Therefore, while the LRP is at stage I, based on cell proliferation activity, one can deduce, how many FCs in the transverse plane gave rise to an LRP. There were cases when progeny of only one FC in transverse plane formed a clone (Fig. 1c). In other cases, one FC produced a clone of marked cells and another two tangentially adjacent pericycle FCs did not form clones but divided (Fig. 4a-a’) suggesting that three FCs in transverse plane participated in LRP formation. Similarly, there was another stage I LRP whose FCs divided once (Fig. 4b), and twice (Fig 4b’,b’’). The entire primordium was composed of a clone of marked cells. As all three FCs produced marked clones one can predict that the progenitor pericycle cell that gave rise to three FCs divided twice tangentially. As this type of division was not described to take place in the differentiation zone, most probable model is that this tangential divisions took place within the root apical meristem of the parent root before cells left the meristem. Indeed, the number of pericycle cells in the transverse plane is increasing within the root apical meristem in the shootward direction (Garay-Arroyo et al., 2013, Fig. S4 therein). Other clones of marked cells found in stage I LRP were formed from four FCs observed in the transverse plane (Fig. 4c-e). Peripheral FCs in transverse plane apparently passed lower number of cell divisions (4d,e). Interestingly, a stage I primordium was found (Fig. 4e) which had three FCs in transverse plane corresponding to longitudinal uni-cellular type and one peripheral FC (right arrow) corresponding to longitudinal bi-cellular type of initiation.

**Fig. 4.**
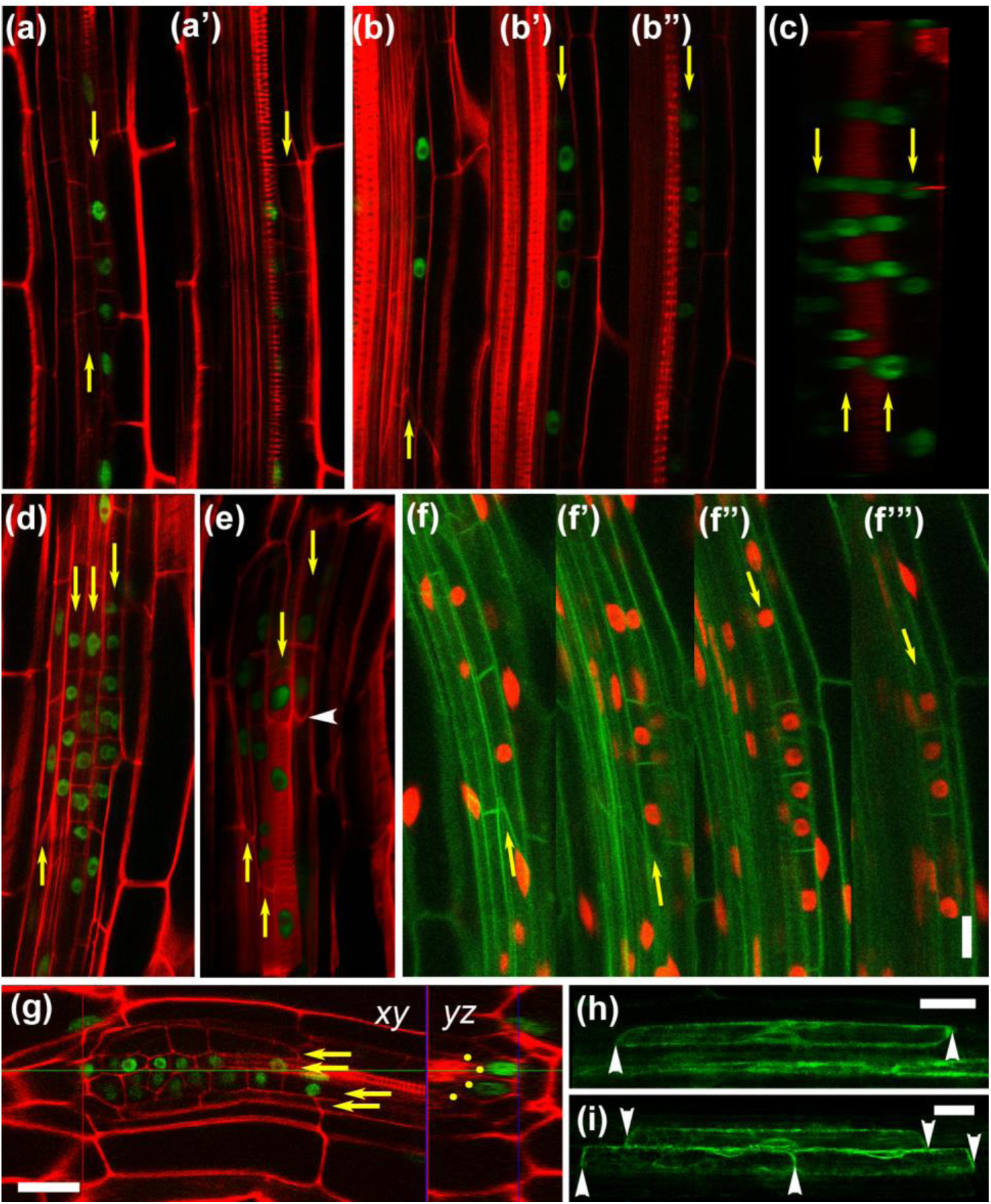
Clones of YFP-marked cells in stage I lateral root primordia (LRPs) and the validation of the number of founder cells (FCs) giving rise to an LRP in the transverse plane. (a, b) LRPs formed from three FCs. One (a) or three (b) FCs in this plane formed clones of YFP-marked cells. (c, d) LRPs formed from four FCs (in the transverse plane), all of which formed a clone of YFP-marked cells. Note that in (d), the peripheral FC on the left underwent one cell cycle (two marked nuclei), whereas other FCs underwent two or more cycles (four to seven marked nuclei). (e) An LRP that formed an asymmetrical clone; three FCs on the left produced clones distributed longitudinally over the entire length of the LRP. The right FC has the same clonal origin, but contributed just half of the LRP extension. An arrowhead indicates the border between the marked and unmarked portions of the LRP. (f) A stage I LRP formed in an F1 seedling from a cross between Wave 131YFP (Geldner et al., 2009) and 35S:H2B:RFP (Federici et al., 2012). (g) An LRP in which two FCs in the transverse plane formed a clone, while two peripheral FCs did not. The yz section shows that, in total, four FCs (circles) gave rise to the LRP. (h and i) An alf4-1;DR5rev:GFP homozygous line in which one (h) and two (i) FCs are specified observed in the root oriented in the protoxylem (h) and protophloem (i) planes. Arrowheads indicate the end walls of the FCs. (c) and (e) are 3D reconstructions, while all other panels show Z-sections in which the FC progeny can be recognised. A cell file corresponding to the progeny of one FC in the transverse plane is indicated by a yellow arrow. Panels labelled with the same letter depict the same primordium. Scale bars: 20 µm. (a–f’’’) are shown at the same magnification.

To validate the conclusion about the number of FCs in transverse plane participating in LRP initiation, early developing LRPs in different genetic backgrounds were analysed. Mostly four FCs in the transverse plane participated in LRP formation (Fig. 4f-f’’’ and Fig. S4). In all cases progeny of one or two FCs can be found at more advanced stages (Cell packets formed from FC on Fig4f’’ and 4f contained 6 and 2 cells, respectively) suggesting that the respective FCs started their activity earlier. In *alf4-1* (*aberrant lateral root formation 4-1*) mutant, deficient in activation of the cell division in pericycle FCs (Celenza et al., 1995; DiDonato et al., 2004; Dubrovsky et al., 2008), one FC in transverse plane is specified more frequently (Fig. 4h,i; number of FCs in transverse plane was 1, 2, and 3 in 62.7, 32.6, and 4.7% of cases, *n* = 43 pre-initiation evens in 11 roots).

Based on clonal analysis of LRPs and emerged LRs, I concluded that most common numbers of FCs participating in LR formation in transverse plane was different in these two age groups. At early stage, deduced number of FCs in transverse plane that gave rise to LRP was 1, 3, 4, and 6 FCs (13.7, 23.5, 37.3, and 2.0 % of all clones found in LRPs, respectively, Table 1). found in emerged LRs, respectively, Table1). This analysis thus suggests that during LR formation, the number of pericycle FCs in this plane is increasing from 1-4 to 6, which in turn indicates that the core (central) FCs can gradually recruit pre-existing neighbour cells to become FCs after LR initiation.

### Clonal analysis reveals that pericycle cells are involved in lateral meristem formation early in root development

When pericycle cells in the mature differentiation zone start to divide periclinally, they give rise to LMs (Dubrovsky and Rost, 2012). In *Arabidopsis* roots, cambium formation starts in relatively young root portion, 15-25 mm from the root tip (Baum et al., 2002). Two types of clones were identified that indicate that pericycle cells are involved in LM formation early in development.

#### Anticlinal pericycle cell divisions in the differentiation zone do not always contribute to LR formation

Clones of two to six relatively short pericycle cells, that did not contribute to LR formation were identified at the xylem pole. Cells were 42 to 98μm long and 5 to 8μm wide (Fig. 5a,b). 26 clones of this type (19%) were identified. In the transverse plane, tangentially, these contained one to three cells (Table 1, Fig. S5) and the respective divisions probably took place within the apical meristem. In some of these clones, after anticlinal divisions pericycle cells additionally passed periclinal ones (Fig. 5c,d). As the cells of these clones were longer than stage I LRP cells (compare Fig. 4 with Fig 5c,d), these clones did not contribute to LRP formation. Periclinal divisions were found in both, relatively young differentiated tissues, where metaxylem elements have not yet completed their differentiation (Fig.5c, 10 clones, Table1), and in more mature root zone (Fig. 5d, 16 clones, Table1).

**Fig. 5.**
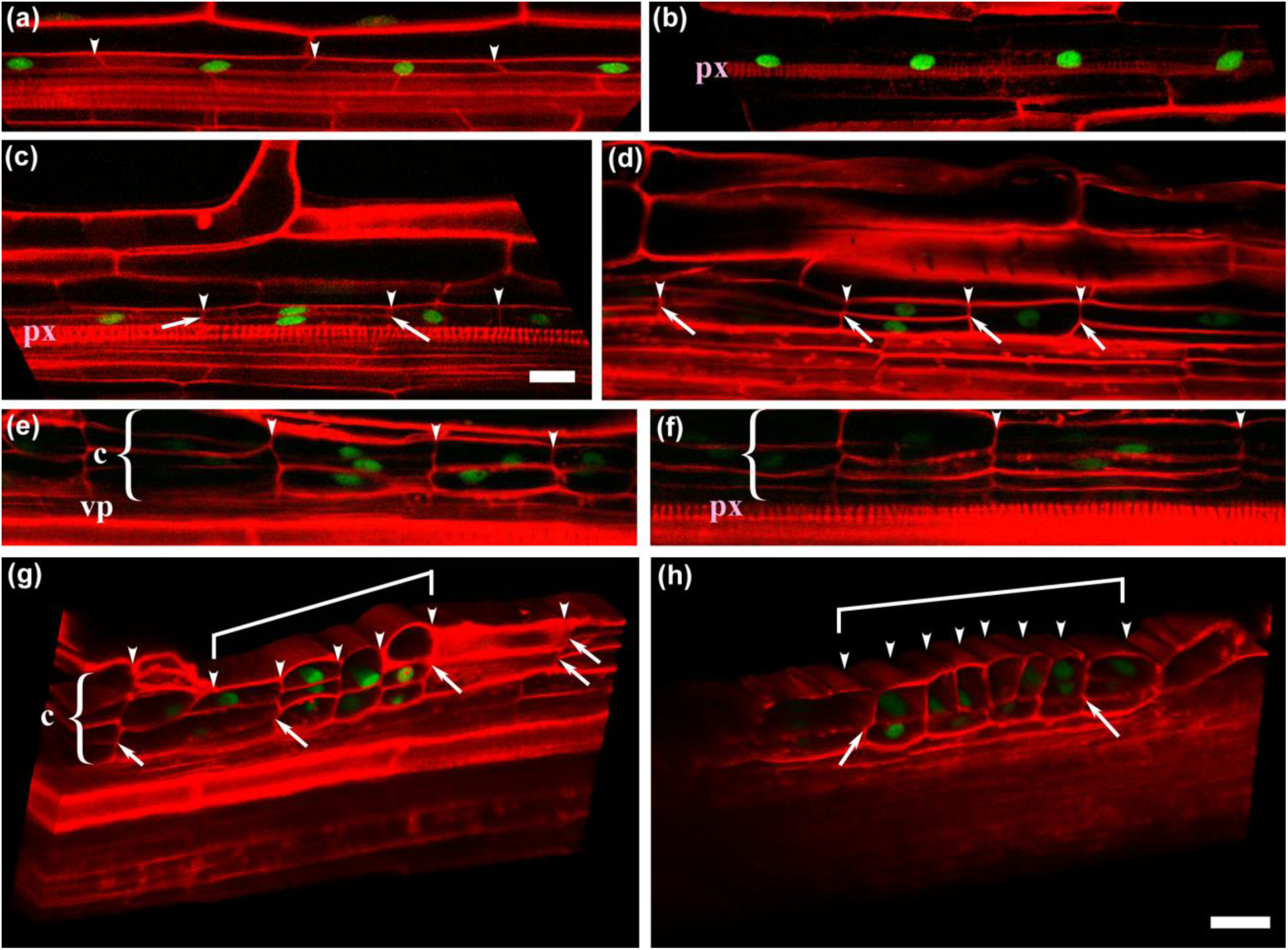
Participation of the pericycle in lateral meristem formation. (a, b) Clones of YFP-marked cells in xylem pole pericycle (XPP) cells. (a) An XPP cell file in a primary root with differentiated protoxylem and metaxylem (not shown). (b) An XPP cell file in a primary root in which only the protoxylem is differentiated. (c, d) Clones of YFP-marked cells formed in XPP cells undergoing anticlinal and periclinal divisions. (c) A clone formed in the relatively young differentiation zone, where only the protoxylem is differentiated. (d) A clone formed in a more mature root portion, where the metaxylem (not shown) is differentiated. (e, f) Clones of XPP-derived cells forming the cambium (c). (e) A clone adjacent to the vascular parenchyma (vp) cells. (f) A clone adjacent to the protoxylem (px) cells. (g,h)Three-dimensional reconstructions of pericycle cell derivatives in the mature root, within a few millimetres of the root base. (g) A clone of YFP-marked cells comprising an arrested lateral root primordium (LRP; square bracket) and adjacent periclinally divided pericycle derivatives comprising the cambium (c, bracket). (h) A similar LRP to (g), containing a greater number of anticlinal divisions. Arrowheads indicate cell walls arising from anticlinal divisions; arrows indicate cell walls resulting from periclinal divisions. Scale bars: 20 µm; (a–c) and (d–h) have the same magnifications.

In the shootward root portion, in the xylem pole, sectors of marked cells arranged in piles of two to three cells (Fig. 5e,f) were found. These arrangements resulted from periclinal divisions of XPP cells. Piles of both marked and unmarked cells contained two to four cells and were contiguous to protoxylem (Fig. 5f) or adjacent to the protoxylem vascular parenchyma (Fig. 5e) cells. Consequently, as these cells were derivatives of pericycle they must be considered the first cambium cells. Pericycle at this stage is not present any more.

In summary, it was found that early in development some pericycle cell divisions form relatively short pericycle cells that in turn undergo longitudinal periclinal division leading to LM formation. When cambium cells are produced (Fig. 5e,f), the length of cambium cells is within the same range as of pericycle cells depicted on Fig. 5a,b. Therefore, anticlinal divisions of XPP cells precede subsequent periclinal divisions. Periclinal and anticlinal divisions found in pericycle in the young differentiation zone unrelated to LR formation suggest that pericycle activity associated with cambium formation can start much earlier than previously considered.

#### The same cell divisions of pericycle in the differentiation zone can simultaneously contribute to lateral root and lateral meristem formation

It has been previously reported that delayed or arrested LRPs can be found between already developed LRs (Dubrovsky et al., 2006; Moreno-Risueno et al., 2010). Within the root portion close to the root base where epidermis, cortex, and endodermis were already lost, clones with cell patterns resembling LRPs (Fig. 5g,h) were found. These clones were formed at the position adjacent to xylem and they were composed of large cells with YFP-marked nuclei arranged in three (Fig. 5h) or four (Fig. 5g) tangentially adjacent longitudinal cell files. These groups of cells probably formed as a result of anticlinal and then periclinal divisions similar to LRPs and contained cells much greater that those of young LRPs initiated in the young differentiated zone (compare with Fig. 3a,c), indicating relatively early (stage III) arrest of their development. Pericycle cells adjacent to these arrested primordia underwent one (Fig. 5h) or two (Fig. 5g) periclinal divisions. At this stage, pericycle itself cannot be recognised anymore and only its derivatives are present. This suggests that pericycle cells in this root portion started periclinal division leading to cambium formation, implying their participation in LM. These data also indicated that arrested LRPs can be a part of the program for the cambium and phellogen formation. Anticlinal divisions within the arrested LRPs can facilitate subsequent periclinal divisions similarly to pericycle anticlinal divisions described in previous section (those depicted on Fig 5C, D)

Pericycle contribution to LR and LM formation are commonly studied separately suggesting these processes are separated in time and space (Esau, 1965; Byrne et al., 1977). If this would be true one could expect to find exclusively clones related to either LM or LR formation. I found, however, clones that included both LRP and LM-related cells. One such clone contained derivatives of the XPP cells (Fig. 6a). Six cells of the clone were located at the base of the developing LRP and the other five cells were located at a flank of the LRP (on Fig. 6a in the lower cell file two nuclei are located outside the shown field of view) and can be considered a part of the developing LR. It is likely that this clone was formed by anticlinal cell divisions of XPP progenitor cell. Pericycle cell file tangentially adjacent to the clone was also composed of short pericycle cells without YFP-marked nuclei (Fig. 6a, a cell file above the clone of marked cells). This clone, therefore, is of a mixed type as it contributes to LRP cells and to relatively short pericycle cells that apparently a LM-related. Similar to clones that contribute to LM formation (Fig. 5a-d), the flanking short pericycle cells should facilitate subsequent periclinal divisions and so one can expect to find clones of mixed type with periclinal divisions. Indeed, part of the same clone formed longitudinally flanking portion near the base of the emerged LR and another, more peripheric part at the LR base, underwent both anticlinal and periclinal divisions (Fig. 6b).

**Fig. 6.**
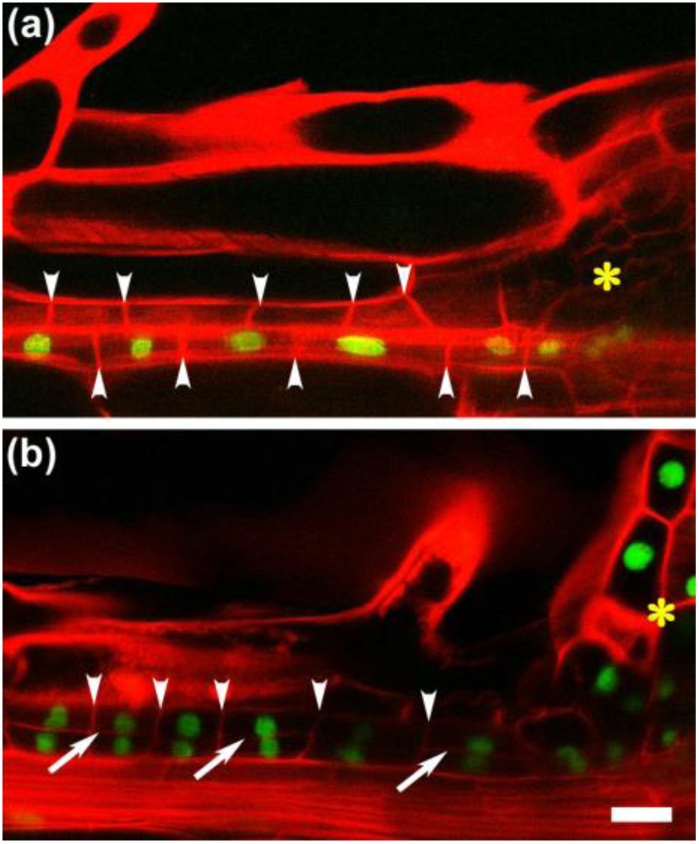
Clones longitudinally adjacent to developing lateral roots (LRs). Clone of YFP-marked and relatively short xylem pole pericycle cells adjacent to developing lateral root primordia (LRP; asterisk). (b) A similar clone to (a), adjacent to a recently emerged LR (asterisk). Note that pericycle cells underwent anticlinal (a) and anticlinal and periclinal divisions. Arrowheads indicate cell walls arising from anticlinal divisions; arrows indicate cell walls resulting from periclinal divisions. Scale bar: 20 µm.

Out of total 39 clones involved in cambium formation (Table 1), 13 clones were found in which one part of the clone constituted the flanking portion of the LR or LRP where anticlinal and periclinal divisions were found. Therefore, one can conclude that pericycle can simultaneously contribute to LRP and cambium formation.

## DISCUSSION

### The pericycle is clonally related to other vascular tissues

There is no consensus on the nature of the pericycle; it is considered either a tissue layer between the ground and vascular tissues (Scheres et al., 1995; Van Norman et al., 2013; Beeckman and De Smet, 2014) or a constituent of the vascular tissue (Parizot et al., 2008; Dubrovsky and Rost, 2012) that is clonally related to the stele (Scheres et al., 1994). Cell-specific microarray data revealed that significantly more genes were simultaneously up-regulated in the pericycle and neighbouring stele tissues in comparison with the number of genes up-regulated in only one of these tissues (Parizot et al., 2012). The clonal analysis presented here clearly demonstrated that, during post-embryonic development, pericycle, protoxylem and vascular parenchyma have a common clonal origin (Fig. 1). This implies that the pericycle and internal stele tissues represent the same tissue complex of provascular tissues in the root apical meristem, which develop from the vascular precursor cells observed at the early globular embryo stage (Wendrich and Weijers, 2013). The discovery that the pericycle is clonally related to other vascular tissues corresponds well with the definition of ‘vascular’ as “plant tissue or region consisting of or giving rise to conducting tissue, xylem and/or phloem” (Evert, 2006, p. 540). Thus, in addition to giving rise to the cambium, which is involved in the formation of secondary conducting tissue, the pericycle and other primary vascular tissue types could have common precursors.

### Stage 0 of LRP formation involves one longitudinal FC undergoing an anticlinal division to produce daughter cells of roughly equal size

It is widely accepted that LR development in *Arabidopsis* starts with three or three pairs of tangentially adjacent FCs (Dubrovsky et al., 2001; Casimiro et al., 2003; Parizot et al., 2008); however, the current analysis revealed that these numbers are more variable. Longitudinally, one to three FCs can give rise to an LRP, but in most cases the main body of the LRP forms from just one cell. Why then have the detailed studies outlined in the Introduction reported that the first initiation events take place in two longitudinally adjacent FCs? I do not see any contradiction here. Inter-primordium distances vary greatly (Dubrovsky et al., 2006), and since the pericycle is an internal tissue, it is very difficult to document the earliest stage when a single FC initiates cell division. This division results in two longitudinally adjacent cells similar to those shown in Fig. 2b (see also Figs. 4b,d (left FC) and 4f (left cell file)). The data show that these two cells are the progeny of one FC, in which *Ds* excision took place. The nuclei of these two cells then migrate, and both cells divide asymmetrically, forming the configuration shown in Fig. 4a (see also Figs. 4Bb, 4d (second FC from the left) and 4f’–f’’). These data indicate that the widely accepted longitudinal bicellular mode of LR initiation is simply a subsequent stage of longitudinal unicellular LR initiation. We must therefore recognise a stage 0 of LRP formation, which implies that the first anticlinal division of the FC results in daughter cells of roughly equal size (as seen in Fig. 4b,d (left cell file) and 4f). Stage 0 LRP would then represent the very beginning of LRP initiation (Fig. 7).

**Fig. 7.**
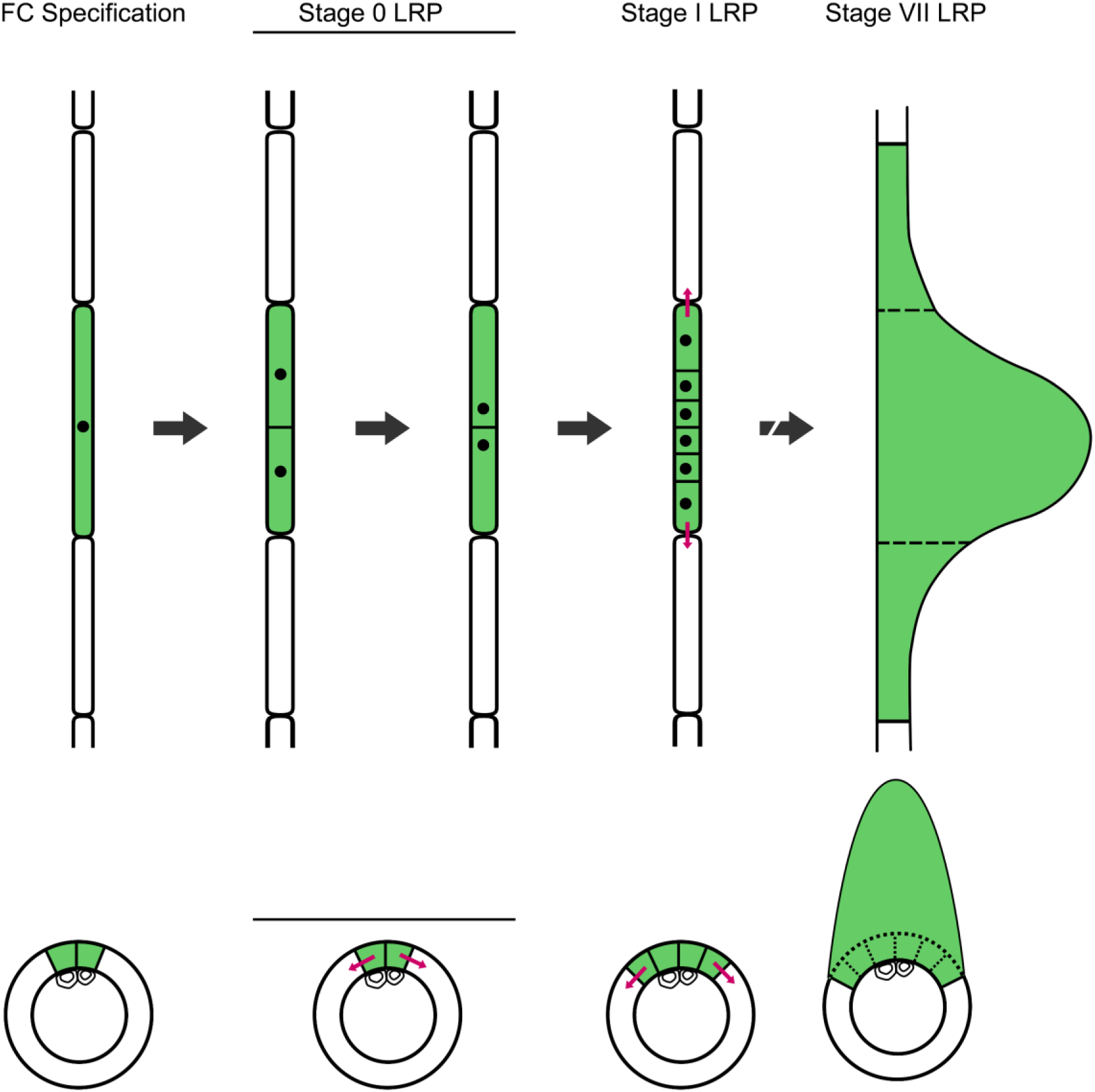
Schematic representation of the gradual involvement of founder cells (FCs) in lateral root (LR) formation. The most common sequence of events is shown. The upper images display the xylem pole pericycle (XPP) cell file in the longitudinal plane. The lower images portray the XPP cells in the transverse plane, with a circle of pericycle cells and protoxylem cells. The first phase shows the specification of the FC (green); two master FCs are specified in the transverse plane. In the stage 0 LRP, FCs undergo their first mitosis and two daughter cells of approximately equal size are formed. At this stage, the nuclei migrate to the newly formed cell wall, resulting in the stage I lateral root primordium (LRP). In the stage I LRP, two more FCs in the transverse plane become recruited to LRP formation (four green cells). During the posterior stages, the longitudinally and transversely flanking cells become involved in LR development, resulting in three cells longitudinally and six cells transversally (marked in green) that participate in LR formation. Red arrows indicate the spread of a signal to recruit new FCs. Black arrows indicate subsequent developmental stages.

### FC recruitment for core and flanking LR regions is asynchronous and FCs have an unequal impact

The clonal analysis revealed that the longitudinal unicellular type of LR initiation was most common. Sussex and colleagues showed that the dimensions of the young LRP correspond roughly to one pericycle cell; however, at later stages when the LR emerges, its longitudinal dimensions correspond to an average of 2.4 pre-existing pericycle cells (Laskowski et al, 1995). This corresponds well with the subset of clones shown in Fig. 6a and b that contribute to the developing LR (Fig. 6). I therefore suggest that the most common scenario during LR formation is the following: (1) longitudinally, a single XPP FC gives rise to an LRP, which becomes the core of the prospective LR; (2) after LRP initiation, two flanking pericycle cells longitudinally and tangentially adjoined to the centrally located FC are recruited and acquire FC identity before LR emergence; and (3) the participation of the later recruited FCs in LR formation becomes more notable after the LR has emerged, and their progeny may be important for the mechanical support of the LR (Fig. 7). The timing of FC recruitment for the core and flanking portions of the LR and their participation in LR morphogenesis suggests their asynchronous involvement, and the unequal participation of XPP cells in the formation of LRs.

An analysis of the participation of the pericycle FCs in the transverse plane revealed that their numbers increase from 1 to 6 during the transition from the early-to late-stage LRP and subsequent LR. In the stage I LRPs, the central (core) FCs give rise to a greater number of derivative cells, implying that one or two centrally located cells behave as master FCs (Wangenheim et al., 2016). Gradual recruitment of FCs confirms the unequal participation of XPP cells in the formation of LRs (Kurup et al., 2005). The most plausible model is that the first FCs initiate LRP formation and recruit the surrounding pericycle cells to become additional FCs, possibly via the auxin-mediated induction involved in master FC recruitment (Dubrovsky et al., 2008). This later recruitment seems to be related to the proliferation of earlier recruited FCs, as no later recruitment is found in the *alf4-1* mutant (Figs. 4H, 4I). This model potentially permits plasticity in LR formation; if no new FCs are recruited, primordium development could arrest or the LRs could be narrower or emerge later in development.

### The pericycle simultaneously participates in the formation of secondary meristems and LRs

The first periclinal divisions leading to the formation of the vascular cambium occur in the vascular parenchyma cells (Baum et al., 2002; Nieminen et al., 2015). The current study showed that these cells divide within the differentiation zone (Fig. 1c) before any other signs of secondary growth can be detected, which suggests that not only the pericycle but also vascular parenchyma cells maintain proliferation outside the apical meristem. These divisions can be considered a preparation for subsequent periclinal divisions that give rise to the residual cambium. Periclinal divisions in the vascular parenchyma cells are followed by a periclinal division in the XPP cells (Baum et al., 2002). Clones resulting from pericycle cell divisions that were unrelated to LRP formation were found in the relatively young differentiation zone (Fig. 5a–c) and were located outside the developmental window for LRP initiation (Dubrovsky et al., 2006). Despite of this we cannot exclude that these relatively short cells are somewhat abnormal LRPs. However, we must disprove this possibility as no abnormal LRPs at later stages were found. Therefore, the presence of these clones indicates that the pericycle cells are preparing to undergo subsequent periclinal divisions. Unexpectedly, periclinal cell divisions in the pericycle were observed in the young differentiation zone (Fig. 5c). Jointly, these observations suggest that the pericycle contributes to the initiation of the LM much earlier than previously thought. The pericycle cells neighbouring arrested primordia undergo a few cycles of periclinal division and form cambium (Figs. 5E, 5F), suggesting that LR formation and the participation of the pericycle in LM formation are co-ordinated. This notion is supported by the clones identified at the longitudinal flanks of the developing primordia or LRs (Fig. 6). In summary, this analysis showed that pericycle participation in LR formation is coupled with its participation in LM formation early in development, and that these activities are not necessarily separated in time and space.

### The immediate pericycle progeny lack self-renewal capacity

Pericycle cells are sometimes considered to be stem cells (Sugimoto et al., 2011; Beeckman and De Smet, 2014). This view emerged because of the general understanding that the pericycle maintains meristematic properties outside the root apical meristem (Dubrovsky et al., 2000; Beeckman et al., 2001) and that it has a high capacity for LR and callus formation and regeneration in vitro (Atta et al., 2009; Sugimoto et al., 2010). By definition, stem cells are those that are simultaneously capable of self-renewal and the production of new cell types (Sugimoto et al., 2011). The clonal analysis performed here showed that no parent root pericycle cells are retained at the sites of FC specification; the FCs enter a new developmental program and their progenies become part of the nascent LRP. Whether pericycle cell identity in the prospective LR is maintained in the FC progeny is an open question, and a previous cell-type-specific gene expression analysis (Malamy and Benfey, 1997) has suggested that this is a possibility. Nevertheless, no self-renewal property is maintained in the immediate XPP progeny within the parent root; the clonal analysis demonstrated that when the XPP cells undergo periclinal division and form the vascular and cork cambia, all pericycle progeny acquire the identity of the LM. Consequently, in strict terms, the pericycle cannot be considered to have a stem-cell identity.

## Acknowledgements

This work was performed during sabbatical study leave of the author in the Department of Plant Sciences, University of Oxford in the laboratory of Liam Dolan. His encouragement, infrastructure and microscopy support, valuable help in the data analysis and their interpretation are gratefully acknowledged. I am also very much grateful to Smita Kurup, Laurent Laplaze, Jim Haseloff, Niko Geldner, John Celenza, Jiři Friml, and Tom Beeckman for seed donations, to Ian Moore and Charlotte Kirchhelle for advice and help on confocal microscopy, Helen Prescott, Marcela Ramírez-Yarza, Selene Napsucialy-Mendivil, Arturo Ocádiz-Ramírez, and Juan Manuel Hurtado-Ramírez for technical help, Natalia Doktor and Gustavo Rodríguez-Alonso for help with art work, and Ari Pekka Mähönen and Liam Dolan for critical reading of previous versions of the manuscript. Sabbatical support was provided to J.G.D. by Mexican Scientific and Technological Council (CONACyT grant 206843) and DGAPA-PASPA-UNAM, Mexico. Research in J.G.D. lab was supported by Mexican CONACyT (237430) and DGAPA-UNAM (IN205315).

## Supporting Information

Additional supporting information includes the following documents.

**Supplementary** Materials and Methods

**Fig. S1.** Root growth dynamics in wild type and *35S-DS1-H2B:YFP; HS-Ac seedlings*.

**Fig. S2.** Anatomical coordinate terminology used in this study.

**Fig. S3.** A lateral root with a clone of marked cells that is spread to approximately one sixth of the progeny of all the FCs analysed in the transverse plane.

**Fig. S4.** Stage I lateral root primordium in *Wave131YFP;pGATA23::NLS-GFP* seedlings.

**Fig. S5.** A clone of pericycle cells that do not contribute to LR formation.

**Table S1.** Technical features of the illustrations presented in the main text and methods of their acquisition

