## Supplementary Materials for "Clonal analysis reveals gradual recruitment of lateral root founder cells and a link between root initiation and cambium formation in *Arabidopsis thaliana*"

**by *Joseph G. Dubrovsky***

### Supplementary Materials and Methods

In most cases clonal analysis of *35S-DS1-H2B:YFP; HS-Ac* seedlings was performed 24 h after a heat shock. We determined if root growth dynamics and developmental pattern in *35S-DS1-H2B:YFP; HS* seedlings is comparable with those in wild type. At the day of heat shock (5 days after germination) in *35S-DS1-H2B:YFP ; HS-Ac* seedlings root length was 71% of that in wild type (Fig. S1). Both wild type and *35S-DS1-H2B:YFP ; HS-Ac* seedlings were subjected to 45 min heat chock. Two days after heat chock, this difference was still present ($P<0.001$, Student's *t*-test) (Fig. S1). LRP development in the*35S-DS1-H2B:YFP; HS-Ac* was indistinguishable from that in wild type. Therefore, we concluded that the developmental pattern in *35S-DS1-H2B:YFP; HS-Ac* seedlings after transposition was similar to that in wild type. Under our growth conditions, the probability of transposition events was low and some plants did not produce clones at all and when they did, frequently a single clone per root was formed. Therefore, we considered similar to Kurup et al. (2005) that that (a) groups of cells with YFP-marked nuclei represent a clone resulting from division of a single mother cell in which transposition took place and (b) the probability of independent and simultaneous transposition events in two neighbouring cells should be low.

Clones previously reported (Kurup et al., 2005, their Fig. 3a, b, d) where longitudinally entire length of the LRP contained YFP-marked cells were interpreted that a single cell in the longitudinal orientation acts as a founder cell to the lateral root as previously defined on the base of histology (Dubrovsky et al., 2001). This interpretation can be opposed by a view that an XPP clone was induced in the primary root meristem and the progenitor cell divided there once or twice and produced a clone of two or four cells in the longitudinal plane. If this would be the case, then after the cells left the meristem, in the differentiation zone two or four longitudinally adjacent YFP-marked cells should be found. Shootward and rootward of the clones depicted on Fig. 2b-f of the main text no long YFP-marked cells were found. Therefore, we have to reject the possibility that the clones were pre-formed in the meristem.

### Supplementary figures

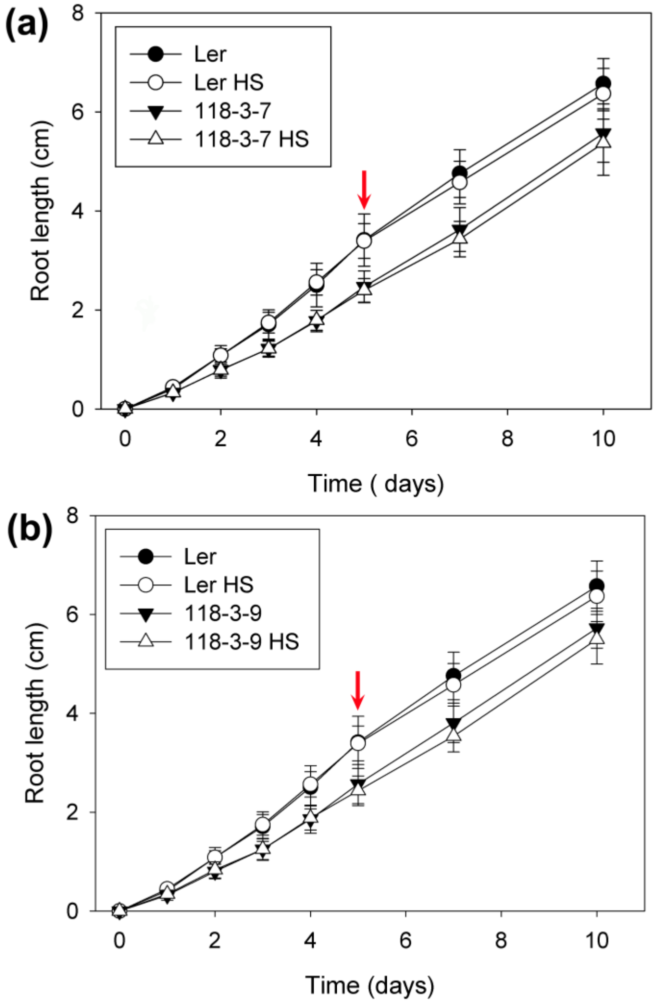


**Fig. S1.** Root growth dynamics in wild type and *35S-DS1-H2B:YFP; HS-Ac seedlings*. Root growth dynamics in F4 generation seedlings. Two sub-lines, 118-3-7 (a) and 119-3-9 (b) were analysed. Red arrows indicate the day when 45 min heat shock treatment was applied. At this day and two days later root length in *35S-DS1-H2B:YFP; HS-Ac* seedling was different ($P$<0.001, Student's $t$-test) compared with the wild type (Ler). HS, heat shock treatment. Data of one representative experiment, $n$ = 10-16; two independent experiments were performed.

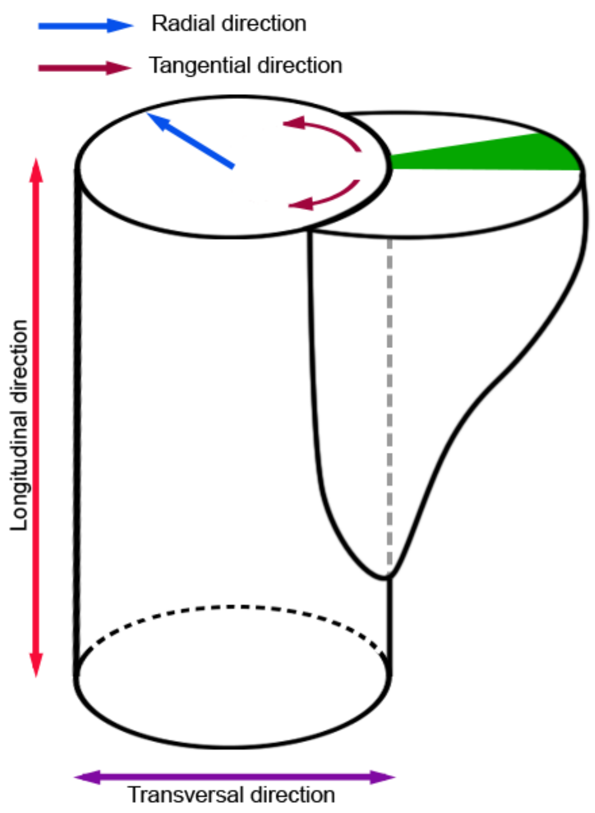


**Fig. S2.** Anatomical coordinate terminology used in this study. Anticlinal cell wall is one perpendicular to the nearest root surface. Periclinal cell wall is one parallel to the nearest root surface.

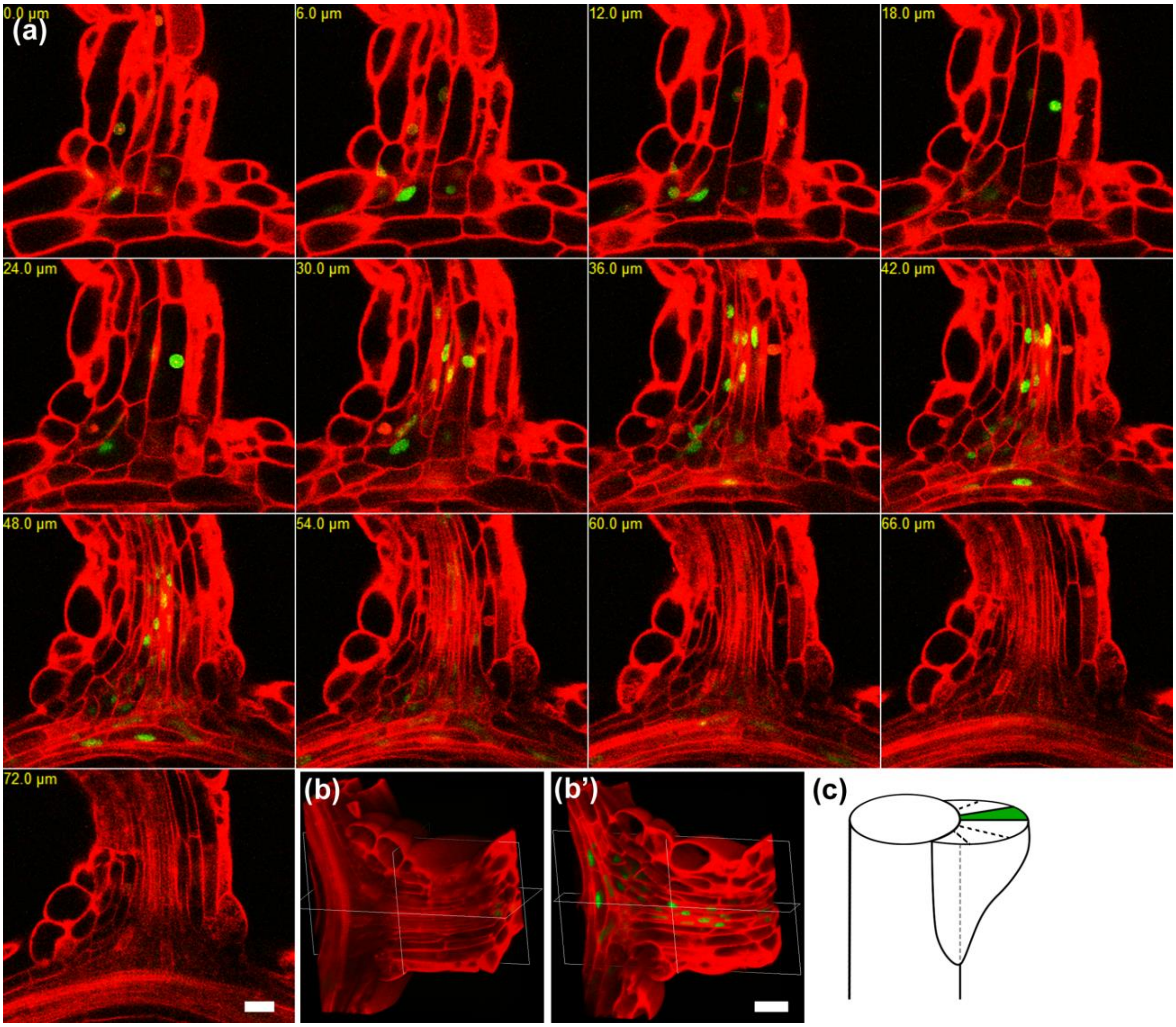


**Fig. S3.** A lateral root with a clone of marked cells that is spread to approximately one sixth of the progeny of all the FCs analysed in the transverse plane. The same LR is shown on Fig. 3E of the main text. (a) Ortho-view of the LR. Note that the first Z-section (0 µm) does not corresponds to Z position where the LR surface begins but is located deeper insight the LR. The deepest section (72 µm) is located at approximately half of the LR thickness. The clone of marked cells occupies approximately 1/3 of the LR portion shown in (a), so it corresponds to approximately 1/6 of the progeny of all the FCs when analysed in the transversal plane. (b) and (b') 3-D reconstruction shows that that the same clone is centrally located when viewed in transverse plane. (c) Interpretation of the same clone. Bars: 20 µm.

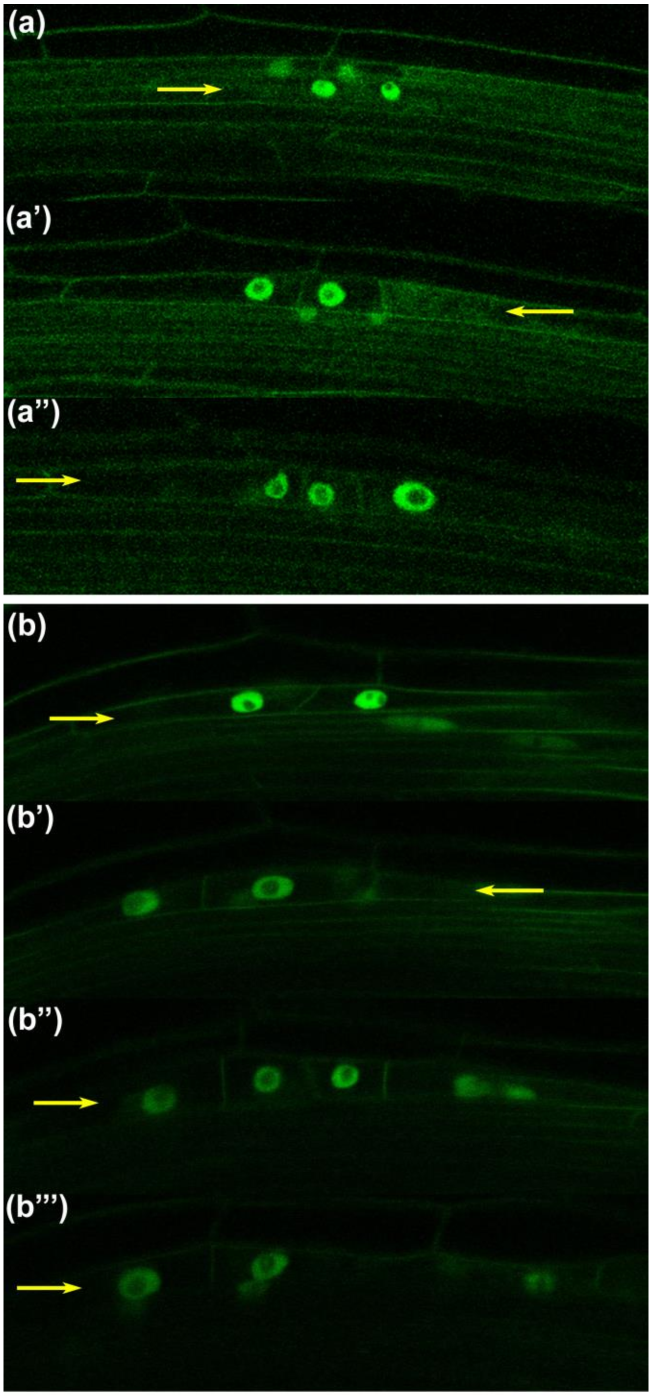


**Fig. S4.** Stage I lateral root primordium in *Wave131YFP;pGATA23::NLS-GFP* seedlings. Analysis of early LRP stages in F1 seedlings of a cross between *pGATA23::NLS-GFP* (De Rybel et al., 2010) and plasmalemma marker line *Wave 131YFP* (Geldner et al., 2009). (a) Z-sections showing that three FCs in transverse plane give rise to a primordium. (b) Z-sections showing that four FCs in transverse plane give rise to a primordium. Each pericycle cell file is indicated with yellow arrow.  This analysis showed that rarely three (11%) and mostly four (89% of total 9 LRPs in 7 roots analysed) FCs in transverse plane give rise to LRP. Bars: 20 µm.

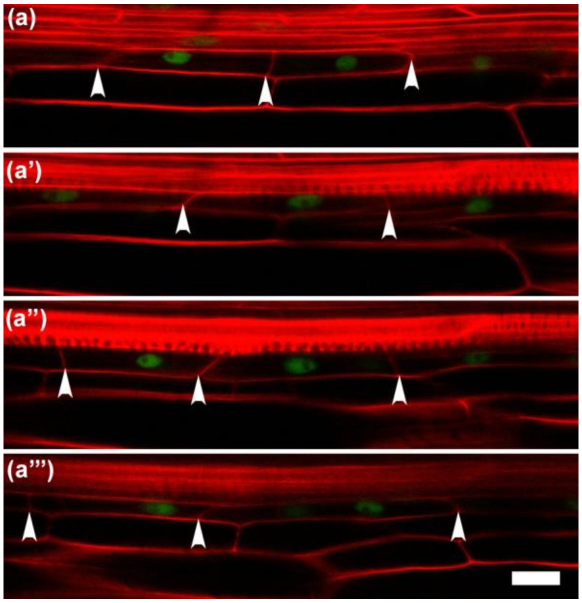


**Fig. S5.** A clone of pericycle cells that do not contribute to LR formation. A clone of tangentially adjacent XPP cells. Each panel corresponds to a different Z sections and shows each of four pericycle cell files involved in the formation of the clone. The clone is located in the mature differentiation zone where metaxylem is clearly recognised (a''). Arrowheads indicate cell walls resulted from anticlinal or oblique divisions. Bar: 20 µm.

**Table S1.** Technical features of the illustrations presented in the main text and methods of their acquisition

| Figure Number | Representation | Enhancement | Optical Sections |
|---|---|---|---|
| 1b, b' | Ortho-view in Zeiss LSM image browser | No | Single section |
| 1c, c' | 3-D reconstruction with MorhoGraphX | No | Single section |
| 1d, d', d'' | Ortho-view in Zeiss LSM image browser | No | Single section |
| 1e | 3-D reconstruction with MorhoGraphX | No | Single section |
| 2a | Maximal intensity projection (Image J) | No | 3 z-sections, 2 µm step |
| 2b | Zeiss LSM image browser | No | Single section |
| 2c | Maximal intensity projection in Z (Image J) | No | 4 z-sections, 2 µm step |
| 2d | Transparent projection mode of Zeiss LSM image browser | No | 7 z-sections, 3 µm step |
| 2e | Standard deviation type of projection in Z (Image J) | No | 21 z-sections, 1.5 µm step |
| 2f | Transparent projection mode of Zeiss LSM image browser | No | 32 z-sections, 2 µm step |
| 2h | Transparent projection mode of Zeiss LSM image browser | No | 29 z-sections, 2.5 µm step |
| 2j | Maximum projection mode of Zeiss LSM image browser | No | 67 z-sections, 2 µm step |
| 2l | Transparent projection mode of Zeiss LSM image browser | No | 24 z-sections, 2 µm step |
| 2m | Transparent projection mode of Zeiss LSM image browser, 3D reconstruction | No | 44 z-sections, 2 µm step |
| 2o | 3-D reconstruction with MorhoGraphX | No | 48 z-sections, 2 µm step |
| 2o' | Transparent projection mode of Zeiss LSM image browser, 3D reconstruction | No | 48 z-sections, 2 µm step |
| 3a, a' | Ortho-view of Zeiss LSM image Browser | Yes, Green Channel | Single sections |
| 3c | 3-D reconstruction with MorhoGraphX | No | 53 z-sections, 1.5 µm step |
| 3c' | Ortho-view in Zeiss LSM image browser | Yes, Green Channel | Single sections |
| 3e | 3-D reconstruction with MorhoGraphX | No | 38 z-sections, 2 µm step |
| 3g | Zeiss LSM image browser | No | Single section |
| 3g', g'', g''' | 3-D reconstruction with MorhoGraphX | No | 98 z-sections, 1.5 µm step |
| 4a, a' | Maximal intensity projection in Z (Image J) | No | 3 z-sections, 1.5 µm step |
| 4b, b'. b'' | Maximal intensity projection in Z (Image J) | No | 4 z-sections, 1.5 µm step |
| 4c | 3-D reconstruction with MorhoGraphX | No | 37 z-sections, 2 µm step |
| 4d | Maximal intensity projection in Z (Image J) | No | 5 z-sections, 1.5 µm step |
| 4e | 3-D reconstruction with MorhoGraphX | No | 33 z-sections, 1.5 µm |

| | | | |
|---|---|---|---|
| | | | step |
| 4f-f''' | Maximal intensity projection in Z (Image J) | No | 3 to 4 z-sections, 1 µm step |
| 4g | Ortho-view of Zeiss LSM image Browser | Yes, Green Channel | Single section |
| 4h | Maximal intensity projection in Z (Image J) | No | 2 z-sections, 1.25 µm |
| 4i | Singe optical section | No | 1 z-section |
| 5a | Transparent projection mode of Zeiss LSM image browser | No | 5 z-sections, 1.5 µm step |
| 5b | Transparent projection mode of Zeiss LSM image browser | No | 10 z-sections, 1.5 µm step |
| 5d | Zeiss LSM image browser | Yes, Green Channel | Single section |
| 5e | Average intensity projection in Z (Image J) | Yes, Green Channel | 3 z-sections, 2 µm step |
| 5f | Average intensity projection in Z (Image J) | Yes, Green Channel | 3 z-sections, 2 µm step |
| 5g | 3-D reconstruction with MorhoGraphX | Yes, Green Channel | 42 z-sections, 1.5 µm step |
| 5h | 3-D reconstruction with MorhoGraphX | Yes, Green and Red channels | 50 z-sections, 2.6 µm step |
| 6a | Maximal intensity projection in Z (Image J) | Yes, Green channel | 7 z-sections, 1.5 µm step |
| 6b | Average intensity projection in Z (Image J) | Yes, Green channel | 6 z-sections, 2 µm step |
| S2-A | Gallery diplay of Zeiss LSM image browser | No | 38 z-sections, 2 µm step |
| S2-B,B' | 3-D reconstruction with MorhoGraphX | No | 38 z-sections, 2 µm step |
| S3-A, A', A'' | Zeiss LSM image browser | No | Single sections |
| S3-B to B''' | Average intensity projection in Z (Image J) | No | 3 z-sections, 1 µm step, for each panel |
| S4 | Average intensity projection in Z (Image J) | No | 3 to 5 z-sections, 1.5 µm step, for each panel |
